# A flexible microfluidic system for single-cell transcriptome profiling elucidates phased transcriptional regulators of cell cycle

**DOI:** 10.1101/2020.01.06.895524

**Authors:** Karen Davey, Daniel Wong, Filip Konopacki, Eugene Kwa, Heike Fiegler, Christopher R. Sibley

**Affiliations:** Department of Medicine, Division of Brain Sciences, Imperial College London, Burlington Danes, London, United Kingdom; Dolomite Bio, Unit 3, Anglian Business Park, Royston, United Kingdom; Institute of Quantitative Biology, Biochemistry and Biotechnology, School of Biological Sciences, 1 George Square, Edinburgh University, EH8 9JZ, United Kingdom

**Keywords:** single cell, drop-seq, droNc-seq, sNuc-seq, regulatory networks, cell cycle

## Abstract

Single cell transcriptome profiling has emerged as a breakthrough technology for the high-resolution understanding of complex cellular systems. Here we report a flexible, cost-effective and user-friendly droplet-based microfluidics system, called the Nadia Instrument, that can allow 3’ mRNA capture of ∼50,000 single cells or individual nuclei in a single run. The precise pressure-based system demonstrates highly reproducible droplet size, low doublet rates and high mRNA capture efficiencies that compare favorably in the field. Moreover, when combined with the Nadia Innovate, the system can be transformed into an adaptable setup that enables use of different buffers and barcoded bead configurations to facilitate diverse applications. Finally, by 3’ mRNA profiling asynchronous human and mouse cells at different phases of the cell cycle, we demonstrate the system’s ability to readily distinguish distinct cell populations and infer underlying transcriptional regulatory networks. Notably this identified multiple transcription factors that had little or no known link to the cell cycle (e.g. DRAP1, ZKSCAN1 and CEBPZ). In summary, the Nadia platform represents a promising and flexible technology for future transcriptomic studies, and other related applications, at cell resolution.

## Introduction

Single cell transcriptome profiling has recently emerged as a breakthrough technology for understanding how cellular heterogeneity contributes to complex biological systems. Indeed, cultured cells, microorganisms, biopsies, blood and other tissues can be rapidly profiled for quantification of gene expression at cell resolution. Among a wealth of notable findings, this has led to the unprecedented discovery of new cell populations such as CFTR-expressing pulmonary ionocytes^1^, new cell subtypes such as the distinct disease-associated microglia found in both mice^2^ and humans^3^, and the single-cell profiling of a whole multicellular organism^4^.

Several technology platforms have been devised for single cell transcriptome profiling that principally differ in amplification method, capture method, scalability and transcriptome coverage (reviewed in ^5^). Methods with lower cell throughput (<10^3^) can provide full transcript coverage permitting analysis of post-transcriptional processing at cell resolution^6–8^. Meanwhile, 3′-digital gene expression (3′-DGE) based technologies focus on the 3’ end of mRNA transcripts to allow a higher throughput (>10^4^) at reduced cost^4, 9–11^. A caveat is that such 3′-DGE methods principally report gene-level rather than isoform-level expression. However, recent adaptations allow membrane-bound proteins to be simultaneously monitored alongside the transcriptome via use of antibody-derived barcoded tags that are captured and concomitantly sequenced^12, 13^.

Relevant to this study, droplet-based single-cell RNA-seq is a popular 3′-DGE method that involves the microfluidics encapsulation of single cells alongside barcoded beads in oil droplets^9, 10^. Cells are subsequently lysed within the droplets and the released polyadenylated RNA captured by oligos coating^9^ or embedded^10^ within the beads for 3′-DGE. Since all oligos associated with a single bead contain the same cellular barcode, an index is provided to the RNA that later reports on its cellular identity during computational analysis. Meanwhile, unique molecular identifier (UMI) sequences within the oligos provide each captured RNA with a transcript barcode such that PCR duplicates can be collapsed following library amplification. Both custom fabricated^9, 10, 14, 15^ and commercial^16, 17^ microfluidics setups have been developed for droplet-based workflows. However, user flexibility of these systems remains limited.

Here we report a new automated and pressure-based microfluidic droplet-based platform, called the Nadia Instrument, that encapsulates up to 8 samples, in parallel, in under 20 minutes. Accordingly, this allows 3’ mRNA capture of ∼50,000 single cells or individual nuclei in a single run. The Nadia Instrument guides users through all relevant steps of the cell encapsulation via an easy-to-use touchscreen interface, whilst it maintains complete flexibility to modify parameters such as droplet size, buffer types, incubation temperatures and bead composition when combined with the Nadia Innovate. We subsequently demonstrate highly reproducible droplet size, low doublet capture rates and high mRNA capture efficiencies relative to alternative technologies. Further, we leverage our high quality datasets to elucidate active transcriptional regulatory networks at different phases of the cell cycle. This revealed transcription factors such as DRAP1, ZKSCAN1 and CEBPZ, among others, that had little or no previous association with distinct phases of the cell cycle. Taken together, the integrity and adaptability of the Nadia platform makes it an attractive and versatile platform for future single cell applications in which fine-tuning of experimental parameters can lead to improved data quality.

## Results

### An open-platform for flexible single-cell microfluidics

Droplet-based single-cell RNA-seq is a scalable and cost-effective method for the simultaneous transcriptome profiling of 100s-1000s of cells. Here we present the flexible, user-friendly and open Nadia platform that facilitates high integrity co-encapsulation of single cells in oil droplets together with barcoded beads (**Figure 1A-C**). Unlike other custom or commercial systems that depend on mechanical injection, the Nadia employs three pressure-driven pumps to deliver smooth and readily manipulated liquid flows of cell suspensions, barcoded beads and oil into the platform’s microfluidics cartridges (**Figure 1B-C**). Successful co-encapsulation of single cells with individual beads subsequently represents the start point for cDNA library preparation. Between 1-8 samples can be processed in parallel on the Nadia due to the flexible configuration of the machines inserted cartridge (**Supplementary figure 1**), whilst incorporated magnetic stir bars and cooling elements ensure samples remain evenly in suspension and temperature controlled throughout. A touch interface guides the user through all essential experimental steps, whilst optional integration of the paired ‘Innovate’ device provides the user with total flexibility to modify all parameters of each run (**Figure 1A**). Accordingly, new protocols can subsequently be rapidly developed, saved and shared for future application by both the user and the wider research community. Further, no wetted parts and disposable cartridges reduce risk of cross-experiment contamination.

**Figure 1:**
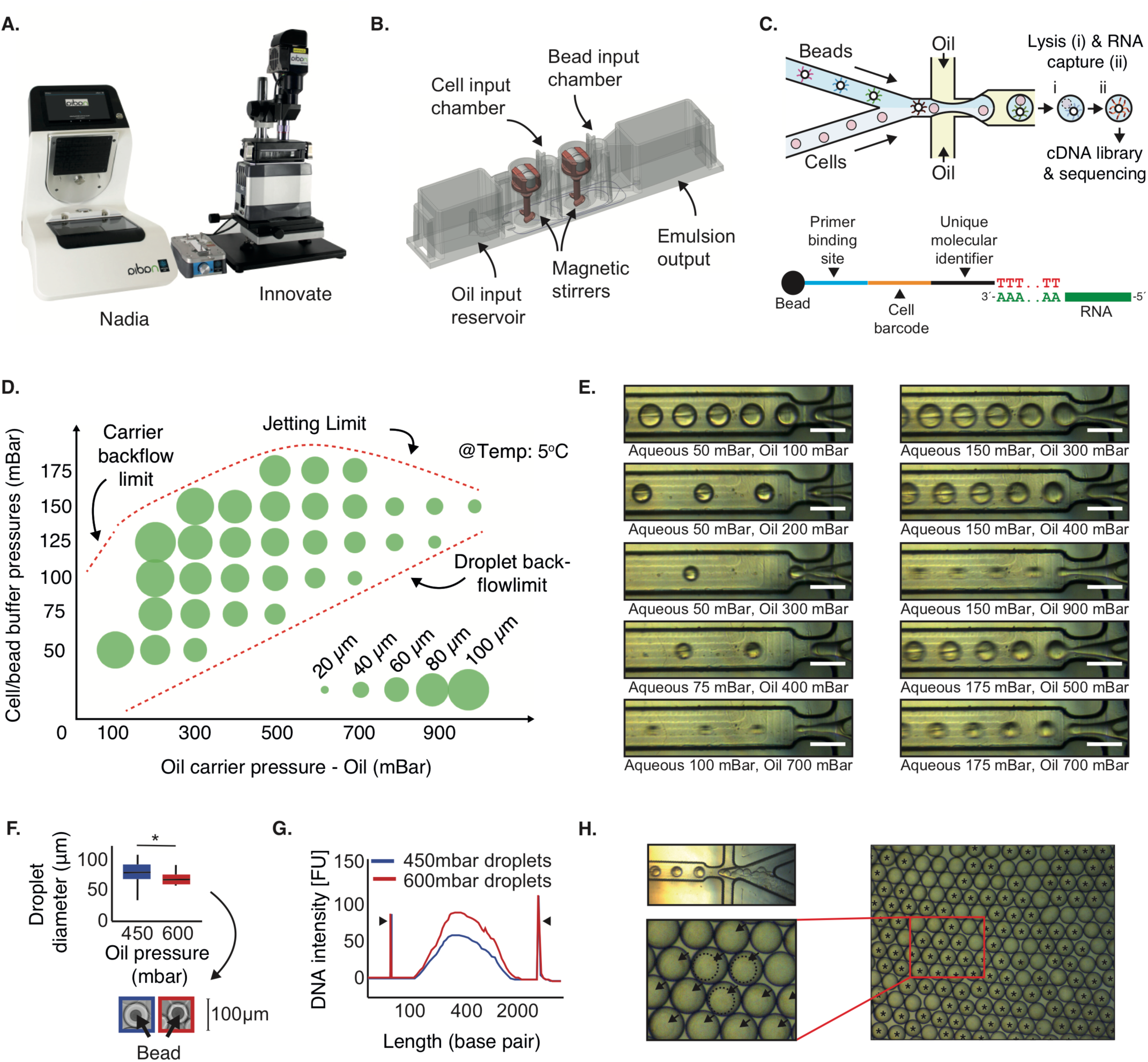
An open platform for single cell transcriptome profiling. **A)** The Nadia Instrument (right) and Nadia Innovate (left) benchtop platform for single-cell transcriptomics. **B)** Design of the disposable microfluidics cartridge used in the Nadia. **C)** Schematic of the droplet sequencing workflow used in the Nadia platform. In brief, single cells or nuclei are encapsulated in oil droplets together with barcoded beads. Following lysis within droplets the released mRNA is captured upon the bead and provided both a cell barcode and a unique molecular identifier. Beads are subsequently pooled prior to reverse-transcription and generation of cDNA libraries called “single-cell transcriptomes attached to microparticles” (STAMPs). The barcoded STAMPs are then amplified in pools for high-throughput RNA-seq. **D)** Theoretical variation of droplet size by changing oil and liquid stream pressures. **E)** Experimental variation of droplet size by changing oil and liquid stream pressures. White scale bars represent 100 μm. **F)** Stable droplet diameters at different oil pressures. Inset shows example droplets containing non-deformable beads. **G)** Bioanalyser traces of full-length transcript PCRs amplified from identical bead numbers but different droplet dimensions. **H)** Example image of deformable beads captured with the Nadia system. Upper left panel shows crowding of deformable beads behind microfluidics junction, lower left panel shows droplet occupancy following sychronised deformable bead loading. For reader guidance, outlines of three deformable beads are indicated with dashed lines, and droplets containing beads are marked by black arrowheads. Right panel shows zoomed out image revealing >70% droplet occupancy of deformable beads. For reader guidance, all droplets containing a deformable bead are marked by a black asterik.

As with related microfluidic setups, single cell suspensions and barcoded beads are loaded at limiting dilutions to ensure minimal occurrence of more than one cell in the same droplet with a bead (**Figure 1C**). Following cell and bead co-encapsulation, the oil droplets act as chambers for cell lysis and mRNA capture. Current injection-based microfluidics systems have been restricted to single droplet sizes^16^, or require custom microfluidics chips designed for purpose^9, 14^. However, retaining the ability to fine-tune droplet volumes could concentrate RNA around oligo bound capture beads for increased mRNA capture, and allow droplet parameters to be optimised according to cell dimensions, buffers or the capture beads used. Exemplifying this, whilst original reports used ∼125 μm diameter droplets for transcriptome profiling whole cells^9^, Habib et al. optimised a microfluidics chip for ∼85 μm diameter droplet generation that facilitated single-nuclei sequencing of archived human brain tissue^14^. Due to the smooth pressure-based system employed, and unlike other platforms, droplet manipulation is readily achieved with the Nadia and accompanying Innovate. Indeed, droplets can be generated over a range of sizes from as little as ∼40 μm (**Figure 1D, E**). Moreover, this can be achieved using the same microfluidics cartridge for all droplet sizes, thus negating the need for custom chip design between experiments. Crucially, resulting droplets are uniform in size (**Figure 1E, F**). Meanwhile, reducing droplet size from ∼85 μm to ∼60 μm (p < 0.05, **Figure 1F**) resulted in increased RNA capture from mouse 3T3 nuclei (**Figure 1G**).

Beyond droplet size control, current droplet-sequencing protocols have principally reported use of two oligonucleotide bound beads; non-deformable beads^9^, and deformable hydrogels^16, 18^. Non-deformable beads have the advantage that mRNA-bound beads can be pooled prior to reverse transcription and minimise reagent costs. In contrast, deformable beads, including those used in commercial platforms^16^, require the reverse transcription reaction to be performed within the droplets to ensure cellular barcodes remain specific to a single cell following oligo release from the hydrogel surface. A reverse transcription mix must thus constitute one of the three streams entering the microfluidics setup which can increase reagent usage. However, whilst droplet-sequencing with non-deformable beads is dependent on double Poisson loading constraints that restricts bead encapsulation to <20%, deformable hydrogels can be efficiently synchronized such that 70-100% of droplets contain a single bead^16, 18^. Whilst the bead configuration is dependent on the application in question, the Nadia importantly retains flexibility to use both non-deformable and deformable beads unlike other platforms^19^. Indeed, whilst non-deformable beads have been used for datasets presented herein, acrylamide/bis-acrylamide deformable beads are fully compatible and allow successful bead stacking behind the microfluidics junction to facilitate synchronised loading of >70% of droplets (**Figure 1H**).

Similar flexibility is provided in the ability to incorporate different buffers. Indeed, stable and mono-dispersed oil droplets are created with a cell/nuclei lysis buffer containing 0.2% sarkosyl and 6 % of the Ficoll PM-400 sucrose-polymer, and a cytoplasmic lysis buffer containing 0.5% Igepal CA-630 (**Supplementary figure 1**). Meanwhile, in an alternative application, use of hyrdogel liquid precursors in replace of the bead-containing lysis buffer can allow hydrogel based capture of the cell suspension to create miniaturized and biocompatible niches for three dimensional *in vitro* cell culture (**Supplementary figure 1**)^20^. Taken together then, the Nadia provides a flexible setup that allows the user to optimise experimental parameters for specific purpose.

### Technical performance for single cell and single nuclei sequencing

In order to test the integrity of the Nadia platform, we performed a mixed-species experiment in which a 3:1 mix of human HEK293 cells and mouse 3T3 cells were subject to droplet capture using the standard machine parameters. During cDNA library preparation, 2000 beads were processed into a final library for sequencing. This number would theoretically equate to profiling of 100 cells under double Poisson loading constraints, and just ∼1.25% of the total cells collected in this run. Following sequencing at >100k reads per cell, our analysis with the Drop-seq tools pipeline^9^ revealed we had collected precisely 100 single cell transcriptomes attached to microparticles (STAMPs). Of these, 75 had mappings primarily to the human genome, and 24 to the mouse genome (**Figure 2A**). Just 1% had mixed mappings that implied capture of more than two mixed species cells during the microfluidics element of the workflow. Meanwhile, each single species cell had a mean of 1.52% reads from the alternative species to imply a low-level of barcode swapping during library preparation. A low doublet capture rate was maintained when the number of beads used for cDNA library preparation was increased, whilst increasing the loading density of cells revealed an increase in doublets consistent with the double Poisson loading of the platform (**Supplementary figure 2**).

**Figure 2:**
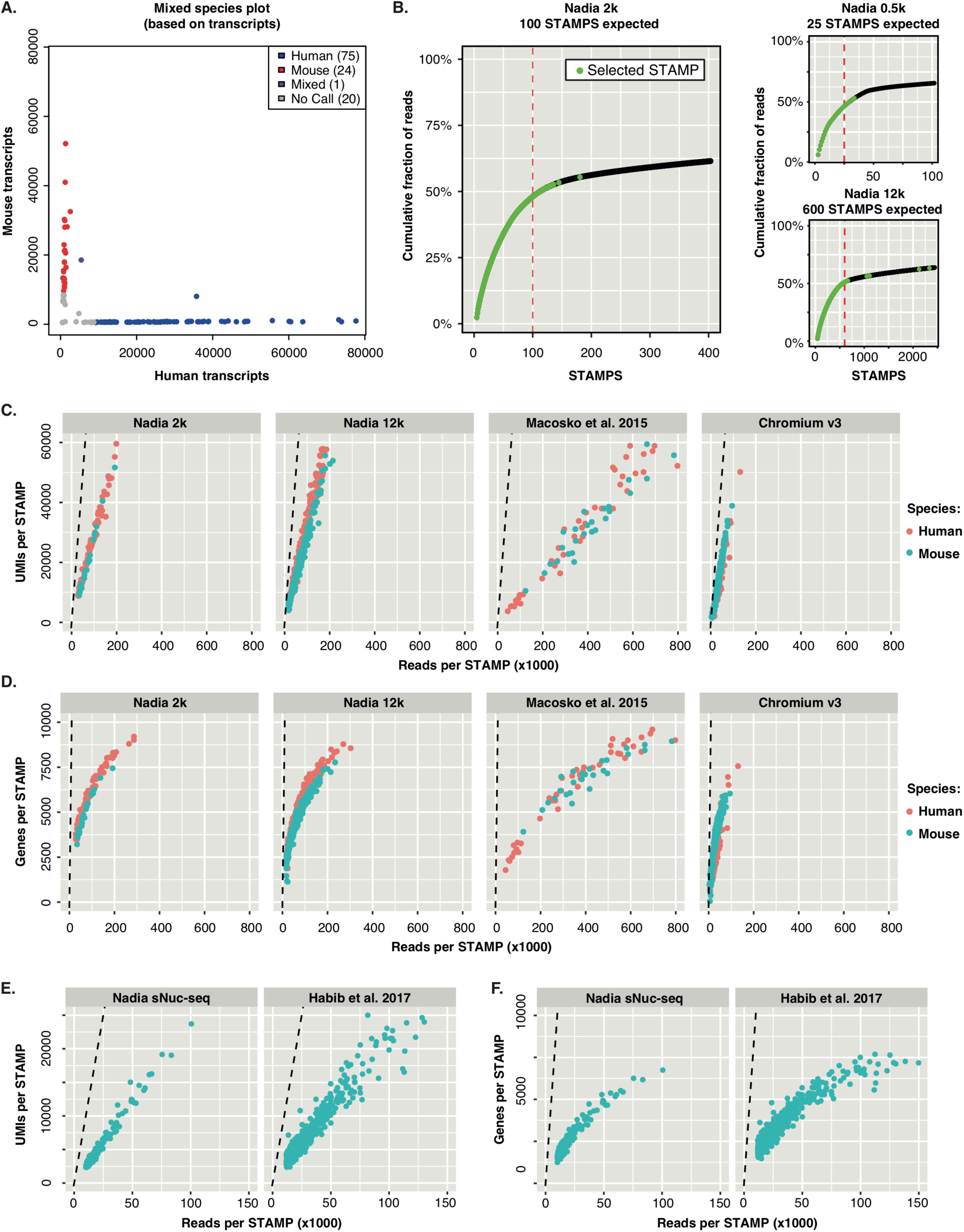
Technical performance for single cell and single nuclei sequencing. **A)** Mixed species barnyard plot of transcripts after profiling 2,000 collected beads (i.e. 100 expected STAMPs) representing a mix of human HEK293 cells and mouse 3T3 cells input at platform recommended cell loading density of 3 x 10^5^ cells per ml. **B)** Cumulative frequency plots reporting sequencing reads associated with individual barcodes when using indicated starting bead inputs for cDNA library construction. Dashed red lines indicate expected STAMPs for each experiment. Larger panel represents dataset used in panel A. “Nadia 2k” generated cDNA libraries from 2,000 beads, “Nadia 0.5k” from 500 beads, and “Nadia 12k” from 12,000 beads. **C)** Number of UMIs detected relative to individual STAMP read counts for indicated mixed-species whole cell experiments (see methods). “Nadia 2k” profiled 2000 collected beads and expected 100 STAMPs, “Nadia 12k” profiled 12000 beads and expected 600 STAMPs, “Macosko et al. 2015” expected 100 STAMPs, “Chromium v3” expected 1400 STAMPs. Dashed line represents maximal point at which each sequencing read would report a unique UMI. **D)** Same as C but with detected genes reported rather than UMIs. **E)** Number of UMIs detected relative to individual STAMP read counts for indicated mouse 3T3 experiments (see methods). Dashed line represents maximal point at which each sequencing read would report a unique UMI. **F)** Same as E but with detected genes reported rather than UMIs.

We next produced cDNA libraries from different amounts of barcoded beads to determine whether STAMP estimates matched the theoretical cell capture of the system. To assess we evaluated the number of UMI counts associated with cell barcodes, and used subsequent graph inflection points to estimate the cells captured. Across multiple experiments performed by independent users at different locations, we saw that the predicted STAMP capture was well matched to expected cell capture (**Figure 2B**). Further, by comparing UMI and gene counts to the total read counts for each library, we found that using the Nadia platform resulted in a high RNA capture efficiency. Indeed this resulted in complex cDNA libraries that had favorable metrics relative to other custom fabricated^9^ (Macosko et al. 2015) and commercial^16^ droplet sequencing platforms for which comparable human HEK293 and mouse 3T3 mixed-species datasets are available (**Figure 2C-D**).

High RNA capture efficiency will be critical for profiling low input material such as single nuclei. Applying such a strategy is necessary when profiling heterogeneous cell samples that cannot be readily dissociated into single cell suspensions (e.g. due to long cellular projections), or when profiling archived samples not robust to freeze-thaw conditions. As such, single-nuclei sequencing is emerging as a method of choice for study of archived human brain tissue^3, 14, 21, 22^. With such future applications in mind, we evaluated the ability of the Nadia platform to profile single nuclei suspensions of mouse 3T3 cells and human HEK293 cells, or mouse 3T3 cells alone. As with whole cell suspensions, mixed-species plots revealed a low doublet rate (**Supplementary figure 2**). In agreement with previous single nuclei sequencing studies^14, 23^, a higher level of intronic reads were reported relative to whole cells (**Supplementary figure 3**). Meanwhile, we found the Nadia platform had nuclear RNA capture rates that compared favourably to limited publically available single nuclei RNA-seq data and approached whole-cell datasets (**Figure 2E-F**)^14^. Whilst capture was marginally reduced relative to whole-cell profiling, the ability to fine-tune droplet dimensions with the Innovate has potential to improve nuclear RNA capture in future (e.g. ^14^). Indeed, we observed an increase in cDNA generated when droplets were reduced from ∼85 μm to ∼60 μm (**Figure 1H**).

Taken together these experiments demonstrate the reliability of the Nadia platform in delivering expected theoretical performance, and the efficiency of the system for both single cell and single nuclei capture.

### Elucidating transcriptional regulatory networks of the cell cycle

To demonstrate the ability of the Nadia platform to distinguish closely related cell populations, we evaluated gene expression profiles linked to cell-cycle progression in 233 human and 277 mouse cells from our “Nadia 12k” mixed-species experiment. Similar to a previous Drop-seq study^9^, and despite the dataset being generated from two asynchronous cell populations, in both species we were able to use gene expression profiles to infer five phases of the cell cycle that matched previous stages of chemically synchronized cells (**Figure 3A**)^24^. This phase assignment was supported by the cycling expression of certain established and novel cell cycle-associated genes, but not housekeeper genes (**Supplementary figure 4**).

**Figure 3:**
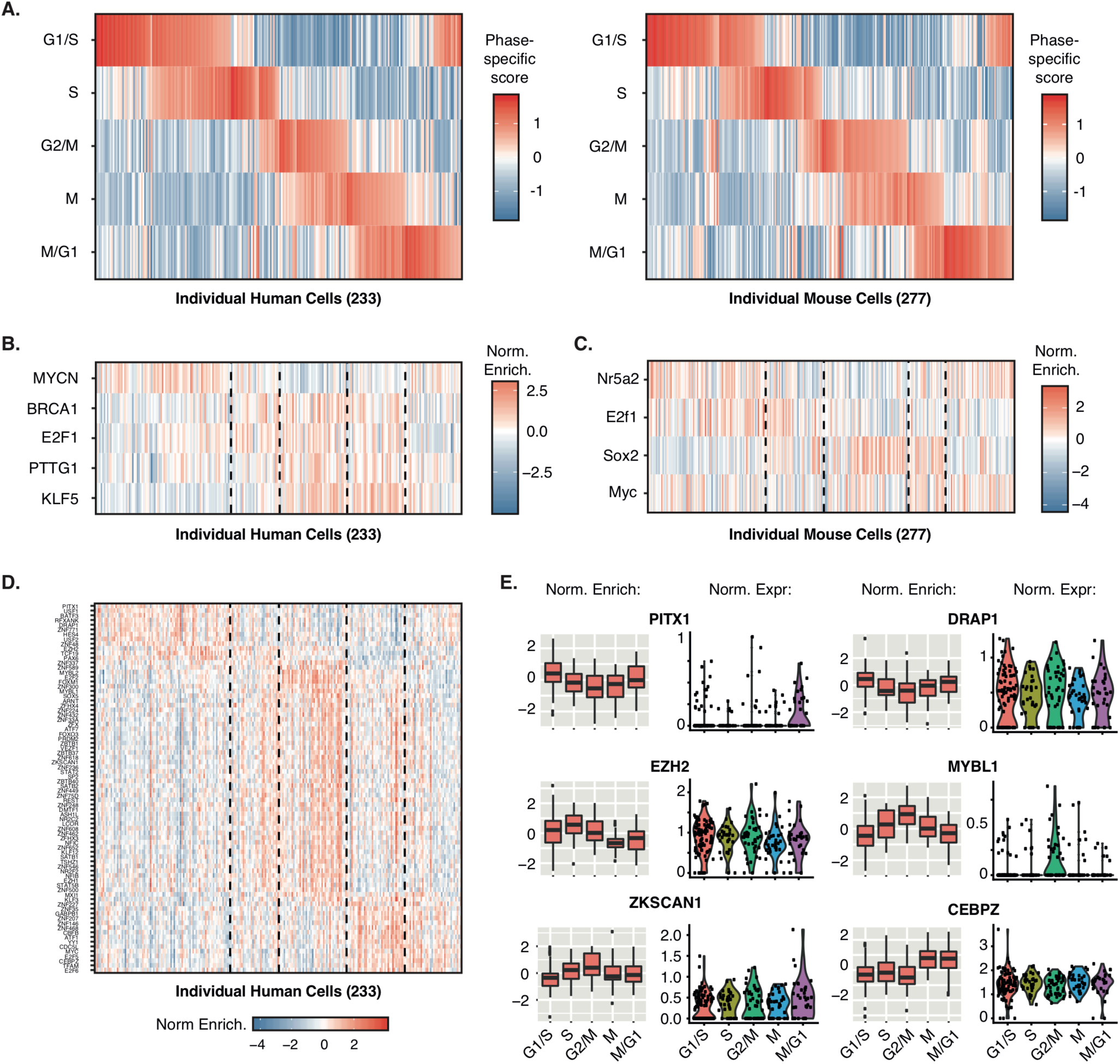
Elucidating transcriptional regulatory networks of the cell cycle. **A)** Inferred cell cycle states of 233 human HEK293 cells (left panel) and 277 mouse 3T3 cells (right panel) based on the gene expression profiles of individual cells relative to stage-specific gene sets (see methods). Cells are ordered by the combination of phases switched on in each individual cell. **B)** Inferred activity of indicated transcription factors based on TRRUST defined regulon expression in individual human HEK293 cells. Dashed lines highlight cell cycle phase assignments used to determine correlation to transcription factor activity. Normalised scores for each transcription factor have been mean centred across all cells. **C)** Same as B but for mouse transcription factors and individual mouse 3T3 cells. **D)** Inferred activity of indicated transcription factors in individual human HEK293 cells based on the summarised expression of regulons that had been inferred from 24 TCGA human cancer tissue sets. Dashed lines highlight cell cycle phase assignments used to determine correlation to transcription factor activity. Normalised scores for each transcription factor have been mean centred across all cells. **E)** Boxplots showing normalised inferred activity and normalised gene expression across different phases for selective transcription factors shown in D. Normalised activity and expression scores for each transcription factor were mean centred across all cells before being summarised by assigned cell cycle phase.

Analysis of single cell gene expression profiles at different stages has previously been used to identify novel genes correlated to cell cycle phases^9^, but the identity of the master regulators that drive coordinated cell-cycle gene-expression programmes remains incompletely understood. Accordingly, we took an alternative approach and questioned whether summarised expression of transcription factor target networks, herein referred to as regulons, could be leveraged to infer the transcriptional regulators active in specific cell cycle phases. Indeed, low depths of sequencing and the absence of mRNA capture for many genes in individual cells (dropouts) can make single cell datasets ineffective in precisely quantitating the expression of individual genes. Meanwhile, many transcription factors can be regulated post-transcriptionally such that their mRNA abundance is not a reliable proxy for protein activity. In contrast, regulon enrichments evaluate differential expression of many transcriptional targets such that these biological and measurement sources of noise are effectively averaged out.

To apply this strategy to the cell cycle we first turned to the manually curated TRRUST database of human and mouse regulons that have been determined from sentence-based text mining^25^. After filtering 800 human and 828 mouse regulons to those expressed in our datasets together with >10 targets, summarised expression profiles were generated for regulons of 77 human and 78 mouse transcription factors across the human and mouse single cells. This revealed select transcriptional regulators whose activity correlated with distinct cell cycle phase scores in both species (p < 0.01, **Figure 3B-C**). Crucially, phase-specific activity aligned with previous studies of these regulators and the cell cycle; KLF5 accelerates mitotic entry and promotes cell proliferation by accelerating G2/M progression^26^, BRCA1 regulates key effectors controlling the G2/M checkpoint^27^, PTTG is active in G2/M phase^28^, MYCN stimulates cell cycle progression by reducing G1 phase^29^, Nr5a2/Lrh-1 knockdown leads to G1 arrest^30, 31^, Myc is a potent inductor of the transition from G1 to S-phase^32^, and Sox2 is a mitotic bookmarking transcription factor active at the M/G1 phase^33^. Notably, E2F1 was found active in G2/M phase of human HEK293 cells and at S-phase of mouse 3T3 cells. This is consistent with its’ control of both G1/S- and G2/M-regulated genes^34^, and E2F1’s role in S-phase progression in mouse 3T3 cells^35^.

Whilst TRRUST reports high confidence and experimentally validated regulons, representation of most transcription factors is limited to few targets. As an alternative, and to further characterise the transcriptional responses of each phase of the human cells in this study, we reasoned regulons inferred by data-driven reverse-engineering methods may offer enhanced opportunity for discovering cell cycle master regulators. Here, VIPER (Virtual Inference of Protein-activity by Enriched Regulon analysis) has recently been developed for the accurate assessment of protein activity from regulon activity^36^, and has recently been extended to single cell analysis via the metaVIPER adaptation^37^. In the absence of previous regulons assembled from HEK293 gene expression profiles, we accordingly evaluated expression of regulons assembled from 24 TCGA human cancer tissue sets using metaVIPER. Indeed, the metaVIPER workflow previously established the utility and integrity of leveraging multiple non-tissue-matched regulons^37^. Encouragingly, this analysis extended our previous findings to reveal a further 78 transcription factors that correlated with one or more phases of cell cycle (p < 0.01, **Figure 3D-E, Supplementary Figure 5**).

Many of the identified transcription factors have previously been identified as master regulators of cell cycle. Among others this included ATF1, SATB2, FOXM1 and MYBL1/B-MYB. Several candidates displayed differential activity in the absence of clear phased-correlated changes in gene expression, thus suggesting activity is regulated by post-translational protein modifications or regulated protein clearance (**Figure 3E, Supplementary Figure 5**). Indeed, only 9/78 were determined as phase-specific genes in previous studies^9, 24^, thus demonstrating the merit of our alternative analysis strategy. Differentially active regulators in the absence of phased gene expression changes included YY1 which is subject to regulatory phosphorylation by various cell cycle associated kinases including Aurora A^38^ and PLK1^39^, FOXM1 that is regulated by SUMOylation^40^ and PLK1 phosphorylation^19^, and REST which is regulated by phosphorylation and USP15 limited polyubiquitination^41^. However, certain transcription factors such as PITX1, SATB2, NR2F2, FOXO3 and MYBL1/B-MYB were regulated at the level of gene expression, likely due to coordinated upstream activity of other master regulators in the cell cycle regulatory gene network.

Last, in addition to known cell-cycle regulated master regulators, we importantly identified multiple differentially active transcription factors that had little or no known link to the cell cycle. This included RFXANK, DRAP1 and HES4 which were correlated with G1/S phase, ZNF33A, VEZF1, ZKSCAN1 which correlated with G2/M phase, and ZNF146, CEBPZ and KLF3 that were maximally correlated with mitosis (**Supplementary Table**). Unlike the others, RFXANK, DRAP1, ZKSCAN1, CEBPZ and KLF3 had no clear relationship between cycling expression levels and activity (**Figure 3E, Supplementary Figure 5**). Accordingly, it will now be important to determine how the phased-activity of these novel cell cycle associated transcription factors manifests in the absence of regulation at the level of gene expression. Indeed, the recent findings that levels of ZKSCAN1 modulate hepatocellular carcinoma progression *in vivo* and *in vitro*^42^, HES4 expression is linked to osteosarcoma prognosis^43^, and that KLF3 loss correlates with aggressive colorectal cancer phenotypes^44^ suggests such understanding could have translational potential. Taken together, our regulon analysis thus confirms, and in several cases extends, understanding of the phase-correlated activity of many transcription factors across cell cycle.

## Discussion

Droplet-based single cell transcriptomics is a more scalable and cost-effective strategy than individual well^45^, FACS^46, 47^ or fluidic circuit-based^48^ alternatives. Here we present a new pressure-controlled and user-friendly microfluidics system that can rapidly enable this powerful strategy to even the inexperienced user. Using pre-fabricated and disposable microfluidics cartridges, the Nadia guides the experimenter through a simple-to-follow workflow that encapsulates ∼8,000 cells per sample, and up to 8 samples in parallel all in under 20 minutes. The paired Innovate add-on provides further opportunity to customise all experimental parameters according to the research question requirements. We present evidence of this experimental adaptability, and report high quality sequencing metrics that compare favourably in the field. We finally demonstrate potential utility of the platform by integrating single-cell transcriptomics with systems biology workflows to extend mechanistic characterisation of the cell cycle. Notably, and among others, we identified DRAP1, ZKSCAN1 and CEPBZ as novel transcription factors with phased-specific activity across G1/S, G2/M and mitosis, respectively.

Flexibility provided by both the Nadia Instrument and the Nadia Innovate is unrivalled by other single-cell microfluidics platforms for droplet based sequencing. Indeed, all parameters of the microfluidics capture process can be modified, including droplet size, stir speeds, incubation temperatures, buffer types and bead composition. The scalability that is achievable through the multiplexed and parallel processing of up to 8 samples can further match or exceed that of other comparable platforms^10, 16^. We demonstrate a high integrity and quality of the transcriptome profiles generated when using the Nadia. Indeed, with standard settings we report a low doublet rate between 1-7% (Fig 2A, Supplementary Figure 2), and favorable RNA capture efficiencies for both single cell and single nuclei sequencing compared to other reports and commercial platforms^9, 14, 16^. Last, the ease-of-use and speed of microfluidics capture will ensure experiment start-to-finish times are kept to a minimum. Accordingly, unintended sample lysis and RNA degradation due to extended protocols is mitigated.

We used the Nadia platform and droplet sequencing workflow to profile the transcriptomes of asynchronous human and mouse cells that subsequently allowed us to infer the different phases of the cell cycle. Notably, the high complexity cDNA libraries allowed us to characterise the cells by transcription factor activity using recently developed systems biology approaches. Our analysis uncovered 83 human transcription factors with inferred activity correlated with one or more cell cycle phase. Despite this, and as noted previously^37^, the employed metaVIPER approach cannot accurately measure activity of proteins whose regulons are not represented adequately in one of the interactomes used for regulon inference. Accordingly, this may explain the absence of overlap between TRRUST curated regulons and those derived from 24 TCGA human cancer tissue sets. However, the expected phase-specific activity of multiple transcription factors (e.g. KLF5, BRCA1, Sox2, Nr5a2, ATF1, SATB2, FOXM1 and MYBL1/B-MYB) when using each source of regulons provides strong support for the validity of the workflow using both sets. The limitation may be mitigated in future as more cell-type specific interactomes are produced.

In addition to confirming phased-activity of many transcription factors such as ATF1, SATB2, FOXM1 and MYBL1/B-MYB, our analysis uncovered several others not previously connected to the cell cycle. This included RFXANK, DRAP1 and HES4 which were correlated with G1/S phase, ZNF33A, VEZF1, ZKSCAN1 which correlated with G2/M phase, and ZNF146, CEBPZ and KLF3. Accordingly, our analysis exemplifies how single-cell transcriptome profiling can be used to further the mechanistic understanding of basic cellular biology. There remains a paucity of knowledge about each of these factors (**Supplementary Table**). It will now be important to experimentally dissect the roles and importance of these novel factors to proliferating cells, how their activity is precisely controlled across phases, and determine their roles in disease. Indeed, the aforementioned links between ZKSCAN1 levels and hepatocellular carcinoma^42^, HES4 levels and osteosarcoma^43^, and KLF3 levels with colorectal cancer^44^ suggests enhanced understanding of these factors in the context of the cell cycle could have translational potential.

In summary then, and as evidenced by our analysis of the cell cycle, the Nadia platforms’s high quality output coupled with its’ flexibility across different buffers, workflows and user-determined parameters suggest it will be an attractive technology for future transcriptomic studies at cell resolution.

## Acknowledgements

This work was supported by an Edmond Lily Safra fellowship and a Sir Henry Dale fellowship jointly funded by the Wellcome Trust and the Royal Society (grant no. 215454/Z/19/Z) to CRS, and funding to CRS from the National Institute for Health Research (NIHR) Biomedical Research Centre based at Imperial College Healthcare NHS Trust and Imperial College London. The views expressed are those of the authors and not necessarily those of the NHS, the NIHR or the Department of Health. All sequencing was performed by the Imperial BRC genomics facility.

## Author contributions

DW, HF and CRS designed experiments. KD and DW performed experiments with contributions from EK, FK and CRS. CRS analysed the data. CRS wrote the manuscript.

## Methods

### Cell preparation

HEK293, HeLa and 3T3 cells were cultured in DMEM with 10% fetal bovine serum (Life Techologies) and 1× penicillin-streptomycin (Life Technologies). Cells were trypsinised for 5 minutes with TrypLE (Life Technologies) before being collected and spun down for 5 min at 300 g. The pellet was resuspended in 1 ml of PBS-BSA (1x PBS, 0.01% BSA) and spun again for 3 min at 300 g. The cells were resuspended in 1 ml of PBS, passed through a 40 μm cell strainer and counted. A concentration of 300 cells/μl in 250 μl of PBS-BSA was subsequently used to allow for the encapsulation of ∼1 cell in every 20 droplets.

### Nuclei suspension preparation

In brief, nuclei isolation media (NIM) was prepared in advance (250mM sucrose, 25 mM KCl, 5 mM MgCl_2_, 10 mM Tris pH8) and pre-chilled. Cells were trypsinised for 5 minutes with TrypLE (Life Technologies) before being collected and spun down for 5 min at 300 g. The pellet was resuspended in 1 ml of PBS-BSA (1x PBS, 0.01% BSA, 0.02 U/μl supernasin) and spun again for 3 min at 300 g. The cells were resuspended in 1 ml of nuclei homogenisation buffer (NIM, 1 μM DTT, 1x Protease inhibitor, 0.1% Triton X-100, 0.04 U/μl RNasin, 0.02 U/μl Superasin) and mixed by gentle pipetting. Sample was then spun at 300g and 4°C for 5 minutes. Supernatant was discarded and the pellet was resuspended in 1 ml of PBS-BSA (0.01% BSA, 0.02 U/μl supernasin). Finally, sample was vortexed and filtered through a 40 μm strainer before nuclei quality was assessed with trypan blue and Hoechst staining and diluted to desired concentration for Nadia loading.

### Microfluidics capture

Cell or nuclei suspensions were captured using the Nadia system according to pre-programmed instrument protocols for drop-seq or sNuc-seq that were accessed through the instruments touch-screen interface. In brief, the Nadia is a fully-automated, bench-top and microfluidic droplet-based platform that can encapsulate up to 8 separate samples in parallel. Each experiment used disposable microfluidic cartridges (covering 1, 2, 4 or 8 samples) with no wetted parts to avoid cross contamination. For each sample, 250 μl of 40 μM-filtered barcoded bead (Chemgene, USA) suspension was loaded into one of the cartridge’s chambers, 250 μl of sample into the second, and 3 ml of oil loaded into the third. Where deformable beads were used, beads were non-barcoded gel beads. Unless specified, cartridge integrated stir bars were set at 75 rpm (cells), 35 rpm (nuclei) and 200 rpm (beads) to ensure that the samples and beads remained in suspension throughout microfluidics capture. Each pre-programmed run lasted 16 minutes and involved bead, sample and oil channels being merged to form oil droplets that co-encapsulated beads together with single cells/nuclei. During each run, three independent pressure pumps controlled the oil, sample and bead channels at pressures up to 1 bar. This ensured consistent conditions and droplet dimensions during each run, whilst providing greatest flexibility to manipulate droplet size and frequency. The standard pressures used were; beads 140 mBar, samples 130 mBar, oil 450 mBar. Double Poisson loading constraints determine that ∼8000 cells/nuclei from a single sample are co-encapsulated with beads when using these default run parameters. Accordingly, 8 samples run in parallel can capture ∼56,000 cells/nuclei during a single run. Additional manipulations of pressure to alter droplet sizes were controlled by the connected Innovate system; an open configurable system used to develop new protocols and applications. Corresponding pressure values are indicated in the text where relevant. Of note, the innovate was connected to a high-speed microscope and camera for real-time droplet formation at the microfluidics junction. Following sample capture in each run, the Nadia’s integrated cooling device was used to chill the samples at 4°C before commencement of library preparation.

### Library preparation

cDNA libraries for 3’ mRNA profiling were prepared using the previously described protocol of Macosko et al. with minor modifications^9^. In summary, mRNA bound beads were removed from the Nadia Instrument’s collection chamber and transferred to a 50 ml falcon tube. Next, 30 mls of 6x SSC buffer (Life Technologies) and 1ml of 1H,1H,2H,2H-Perfluorooctan-1-ol (Sigma Aldrich) were added before mixing via inversion. After spinning at 1000g for 2 minutes the supernatant was removed and retained in a separate falcon whilst being careful not to disturb the beads at the oil-water interface. A further 30 mls of 6x SSC buffer were added to the original sample to disturb the beads before mixing via inversion. Oil was allowed to settle to the bottom before bead containing suspension was transferred to a new falcon tube. After disturbing the oil fraction with a 1 ml pipette to collect any missed beads, both falcons containing ∼30 mls of bead containing SSC buffer were spun at 1000g and 4°C for 2 minutes. At this stage, ∼26mls of supernatant was carefully removed from each tube whilst being careful not to disturb the beads. Beads were subsequently resuspended with retained buffer and transferred to a 1.5 ml eppendorf. Beads were spun down in a desktop micro-centrifuge and buffer removed. Additional bead fractions were added and the process repeated until all beads were collected. At this stage the buffer was removed and all beads washed by pipetting in 1 ml of 6x SSC buffer. Buffer was removed and beads were subsequently washed in 200 μl of 5x Maxima RT buffer (Life Technologies).

Reverse transcription was performed in 200 μl of a 1x RT mix (80 μl nuclease free water, 40 μl of 5x Maxima RT buffer, 40 μl of 20% Ficoll PM-400, 20 μl of 10 mM dNTP mix, 5 μl of RNasin, 10 μl of Maxima H-RT enzyme, 5 μl of 100 μM TSO-RT primer) with the following conditions; 30 minutes at 23°C, 2 hours at 42°C. Throughout the process the sample was set to shake at 1100 rpm. Beads were subsequently spun down, RT mix removed and the beads washed in once in TE-SDS buffer (10 mM Tris pH 8, 1 mM EDTA, 0.5% SDS), twice in TE-TW buffer (10 mM Tris pH 8, 1 mM EDTA, 0.01% Tween-20) and once in 300 μl of 10 mM Tris pH 8. Beads were subsequently incubated for 45 minutes at 37°C and 1100 rpm in Exonuclease I mix (170 μl nuclease free water, 20 μl 10x Exonuclease I buffer, 10 μl Exonuclease I - Life Technologies). Beads were then washed once in TE-SDS buffer, twice in TE-TW buffer and then re-suspended in 300 μl of nuclease free water. Beads were subsequently counted with a haemocytometer after mixing 20 μl of beads with 20 μl of 20% PEG400 (Sigma Aldrich). An average of 4 counts were taken before test PCRs at different cycle numbers were performed with desired bead aliquots for each experiment (∼2000-5000) to gauge optimal cycles for final PCRs on subsequent beads. Specifically, PCR mix included 24.6 μl of nuclease free water, 0.4 μl of 100 μM TSO-PCR primer, and 25 μl of Kapa HiFi readymix (Roche Diagnostics). Cycling conditions were 95°C for 3 minutes, four cycles of 98°C for 20 seconds, 65°C for 45 seconds, 72°C for 3 minutes, followed by variable cycles (∼9-14) of 98°C for 20 seconds, 67°C for 20 seconds, 72°C for 3 minutes. A final extension of 72°C for 3 minutes completed the PCR. At the end of elongation steps during the first four cycles, PCR tubes were removed from machine and beads suspended by gentle agitation.

Following optimised PCRs of desired bead numbers, we enriched cDNA products longer than 300 base pairs using select-a-size spin columns (Zymogen) according to the manufacturer protocol. After bioanalyser evaluation and quantification of products, 550 pg of DNA was used as input for an Illumina Nextera tagmentation reaction according to manufacturer’s protocol (15 μl Nextera PCR mastermix, 8 μl nuclease free water, 1 μl of 10 μM TSO-hybrid oligo, 1 μl of 10 μM Nextera N70X indexed oligo). This reaction reduced cDNA libraries to a size distribution suitable for Illumina sequencing, and added a common PCR handle for 12 cycles of final library amplification (95°C for 30 seconds, twelve cycles of 95°C for 10 seconds, 55°C for 30 seconds, 72°C for 30 seconds, final extension of 72°C for 3 minutes). Last, as shorter cDNA inserts are more likely to be overlap variable length poly-A tails, we again enriched for cDNA products longer than 300 base pairs using select-a-size spin columns (Zymogen) according to the manufacturer protocol. The final library profiles were then evaluated and quantified with a Bioanalyser, Qubit and Tapestation prior to sequencing.

### Next generation sequencing

All high throughput sequencing was performed using an Illumina NextSeq 500 sequencer at the Imperial BRC genomics facility. Samples were run using a custom read 1 primer (Read1customSeq). Read 1 was set at >20 base pairs to read through the cellular and molecular barcodes, and read 2 set at >25 base pairs to read cDNA inserts. Additional 8 base pair index reads were used to determine libraries within multiplexed runs. Each run had 5-10% PhiX spiked in to the library to ensure suitable complexity at low diversity sequencing cycles.

### Oligonucleotides

The following oligonucleotides were used for library preparation and sequencing:

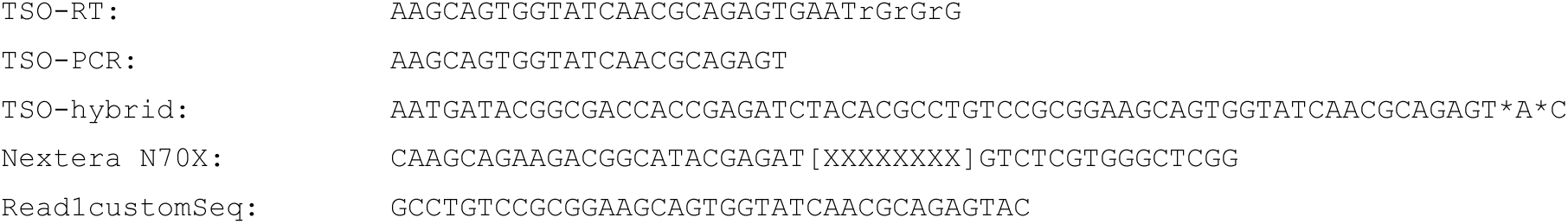

### Data processing

Raw fastq files were processed with the Drop-seq toolkit established in Macosko et al.^9^ according to recommended guidelines. The pipeline was implemented via the DropSeqPipe v0.4 workflow^49^. In brief, Cutadapt v1.16 was used for adapter trimming, with trimming and filtering was performed on both fastq files separately. STAR v2.5.3 was used for mapping to annotation release v.94 and genome build v.38 for Mus musculus, or annotation release v.91 and genome build v.38 for Homo sapiens. Multimapped reads were discarded. Dropseq_tools v2 was used for demultiplexing and file manipulation according to recommended guidelines, and technology-specific positions of the cell barcodes and unique molecular identifiers (UMI) were used. A whitelist of cells barcodes with minimum distance of 3 bases was used. Cell barcodes and UMI with a hamming distance of 1 and 2 respectively were corrected.

For cell cycle phase determination, gene expression profiles of individual cells were related to adapted gene sets used in Macosko et al. that represent distinct phases of the cell cycle^9^. Specifically, phase scores for each cell-cycle stage were determined for individual cells by averaging the log normalised expression levels, derived using Seurat (v3.1.1)^50^, of the genes in each gene-set. The mean scores for each phase were then mean centred and standard deviation normalised across all cells, before phases for each individual cell were mean centred and standard deviation normalised. Cells were subsequently ordered according to the combination of phases determined to be switched on in each individual cell.

### Regulatory transcriptional networks

Datasets were initially filtered to those genes expressed in at least 10% of cells of each single-cell library. Raw counts were subsequently log normalised and scaled with Seurat. For figures 3B and 3C, human and mouse transcription factor targets were downloaded from the TRRUST v2 database^25^. Regulons were subsequently filtered to those expressed in respective human and mouse cell datasets alongside >10 identified targets. Transcription factor activity was subsequently scored in individual cells by averaging the normalised expression levels of the genes in each regulon. The mean scores for each regulon were mean centred and standard deviation normalised across all cells. Normalised inferred regulon activity of individual cells was subsequently correlated with the previously inferred phase-specific scores, with those having a significant (p < 0.01) pearson correlation of >0.3 with one or more phases being used for presentation. The cor.test function of the stats (v. 3.6.1) R package was used for calculation of pearson correlation and test statistics. For figures 3D and 3E, regulons used were previously derived from 24 TCGA human cancer RNA-seq datasets and accessed from the ‘aracne.networks’ R package. VIPER (v.1.18.1)^36, 37^ was used to score all regulons from the 24 TCGA human cancers in all individual human HEK293 cells, before the average of all normalised enrichment scores (i.e. avgScore) for each specific master regulator was used to integrate scores into a single metric. The mean scores for each regulon were mean centred and standard deviation normalised across all cells. Inferred regulon activity of individual cells was subsequently correlated with the previously inferred phase-specific scores, with those having a pearson correlation of >0.35 with one or more phases being used for presentation.

### Data availability

Study generated transcriptomic data has been deposited in the GEO repository and will be made available upon publication. External datasets were collected from following sources: Macasko et al. 2015 mixed species from GEO accession GSE63473 (SRR1748412), Chromium v3 from 10x Genomics (https://www.10xgenomics.com), Habib et al. 2017 mouse 3T3 nuclei from the Broad Institutes Single Cell portal (https://portals.broadinstitute.org/single_cell).

## Supplementary figure legends

**Supplementary figure 1:**
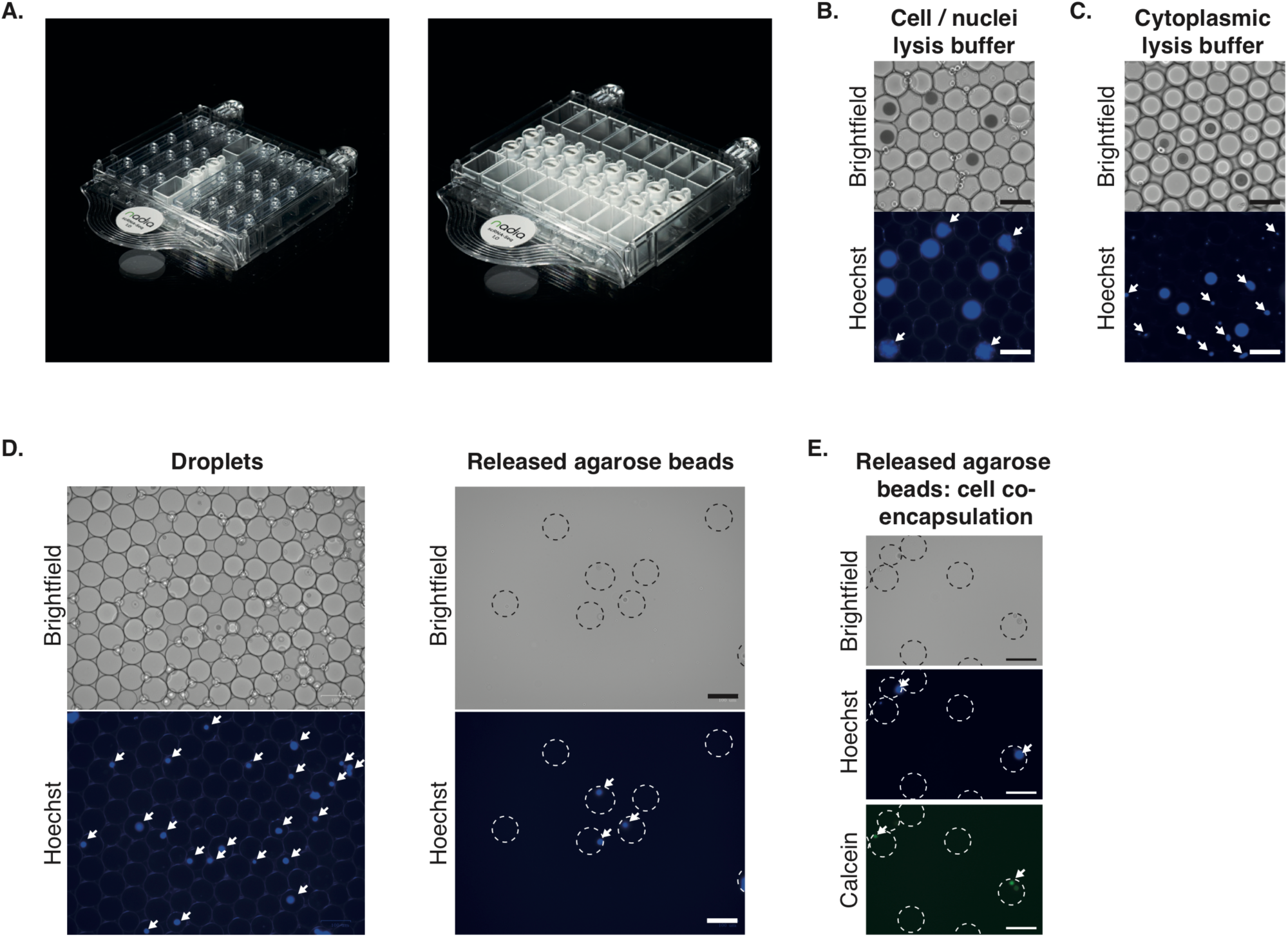
**A)** Nadia cartridge in both 1 and 8 individual microfluidic chip formats. **B)** Nadia generated oil droplets using a cell/nuclei lysis buffer containing 0.2% sarkosyl and 6 % of the Ficoll PM-400 sucrose-polymer. Brightfield shows mono-dispersed droplets and encapsulation of non-deformable beads. Hoechst staining reveals additional droplets where whole cells have been encapsulated and lysed (white arrows). **C)** Nadia generated oil droplets using a cytoplasmic lysis buffer containing 0.5% Igepal CA-630. Brightfield shows mono-dispersed droplets and encapsulation of non-deformable beads. Hoechst staining reveals additional droplets where whole cells have been encapsulated and the unlysed nuclei are stained (white arrows). **D)** Replacement of lysis buffer with hyrdogel liquid precursors (e.g. 1% agarose) allows whole cell microencapsulation. Left panel: Brightfield and imaging of Hoechst-stained HEK293 cells reveals that individual cells were successfully encapsulated at a distribution of ∼1 cell per 5 droplets. White arrowheads indicate encapsulated cells. Right panel: Brightfield and imaging Hoechst-stained HEK cells reveals agarose beads containing HEK293 cells were successfully extracted from the emulsion using perfluorooctanol. White arrowheads indicate encapsulated cells. All agarose beads have been outlined in hashed white lines to aid visualisation. **E)** Same as D except mixed cell populations have been co-encapsulated, and only released agarose beads are shown. Human HEK293 cells are Hoechst stained (middle panel), mouse 3T3 cells have been stained with calcein (lower panel). White arrowheads indicate encapsulated cells. All agarose beads have been outlined in hashed white lines to aid visualisation.

**Supplementary figure 2:**
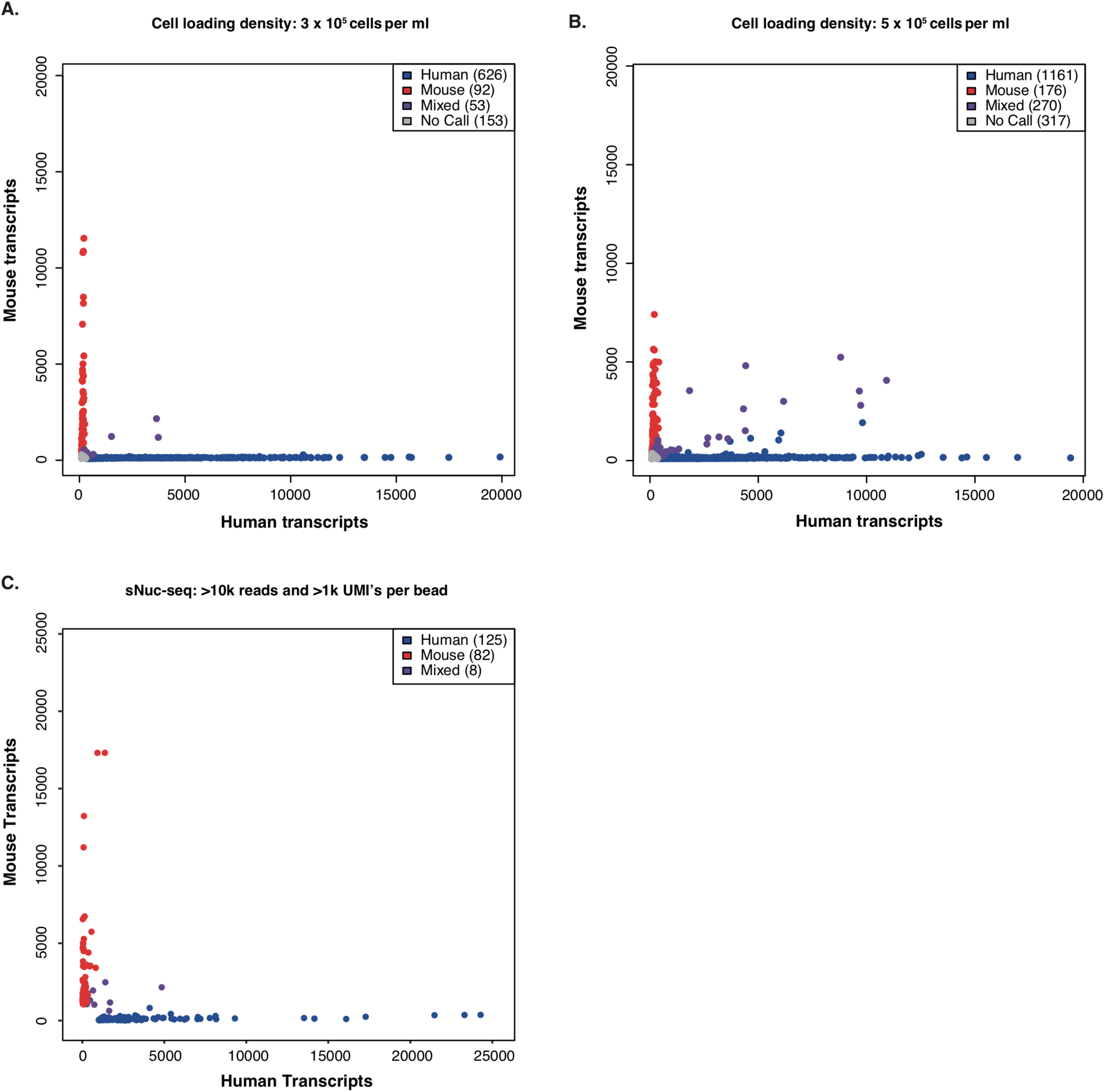
**A)** Mixed species barnyard plot of transcripts after profiling 16,000 collected beads representing a mix of human HeLa cells and mouse 3T3 cells input at platform recommended cell loading density of 3 x 10^5^ cells per ml. **B)** Mixed species barnyard plot of transcripts after profiling 12,000 collected beads representing a mix of human HeLa cells and mouse 3T3 cells input at cell loading density of 5 x 10^5^ cells per ml. **C)** Mixed species barnyard plot of transcripts after profiling a mix of human HEK293 cells and mouse 3T3 nuclei at platform recommended loading densities. STAMPS with less than 1,000 UMIs were filtered out.

**Supplementary figure 3:**
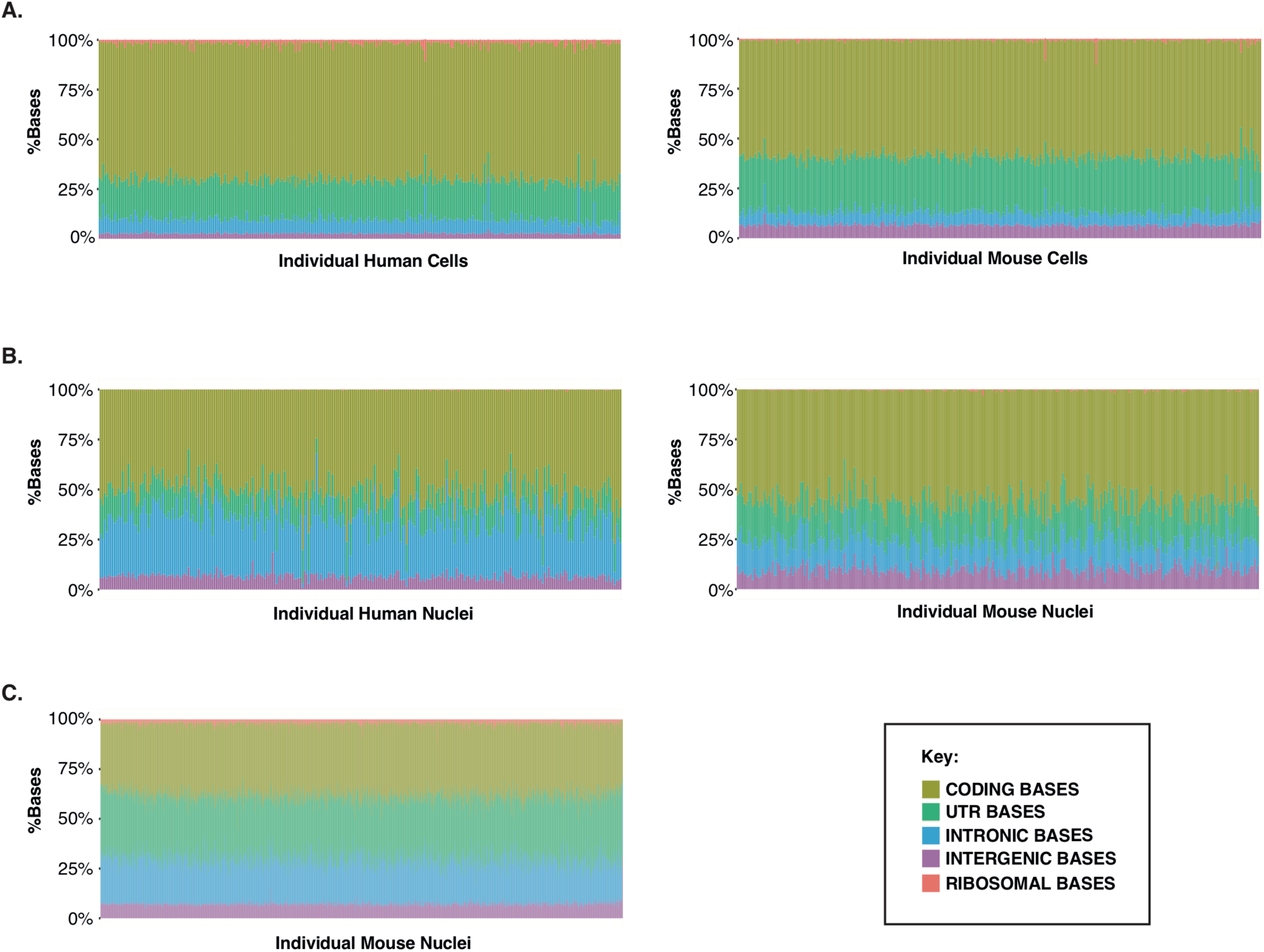
**A)** Percentages of reads mapped to the indicated regions of the human genome for human HEK293 cells (left panel), and percentages of reads mapped to the indicated regions of the mouse genome for mouse 3T3 cells (right panel). Cells detailed are those profiled in Figure 2A. **B)** Percentages of reads mapped to the indicated regions of the human genome for human HEK293 nuclei (left panel), and percentages of reads mapped to the indicated regions of the mouse genome for mouse 3T3 nuclei (right panel). Cells detailed are those profiled in Supplementary Figure 2C. **C)** Percentages of reads mapped to the indicated regions of the mouse genome for mouse 3T3 nuclei. Cells detailed are those profiled in Figure 2E-F.

**Supplementary figure 4:**
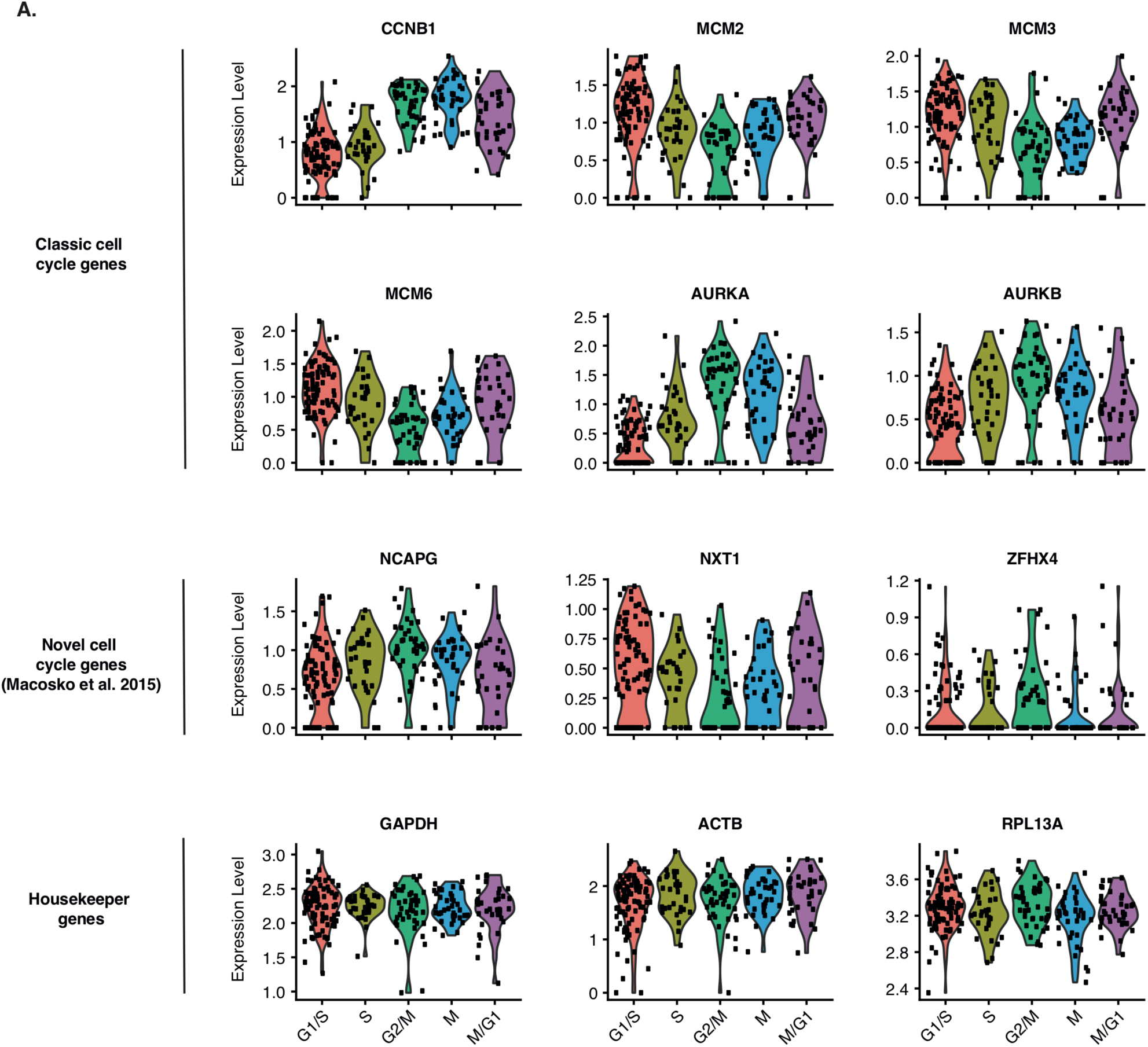
Normalised gene expression profiles of indicated genes across the cell cycle. Shown are classical cell cycle associated genes (top two rows), novel cell cycle associated genes discovered in Macasko et al. 2015 (third row), and housekeeper genes not expected to be correlated with distinct cell cycle phases (fourth row).

**Supplementary figure 5:**
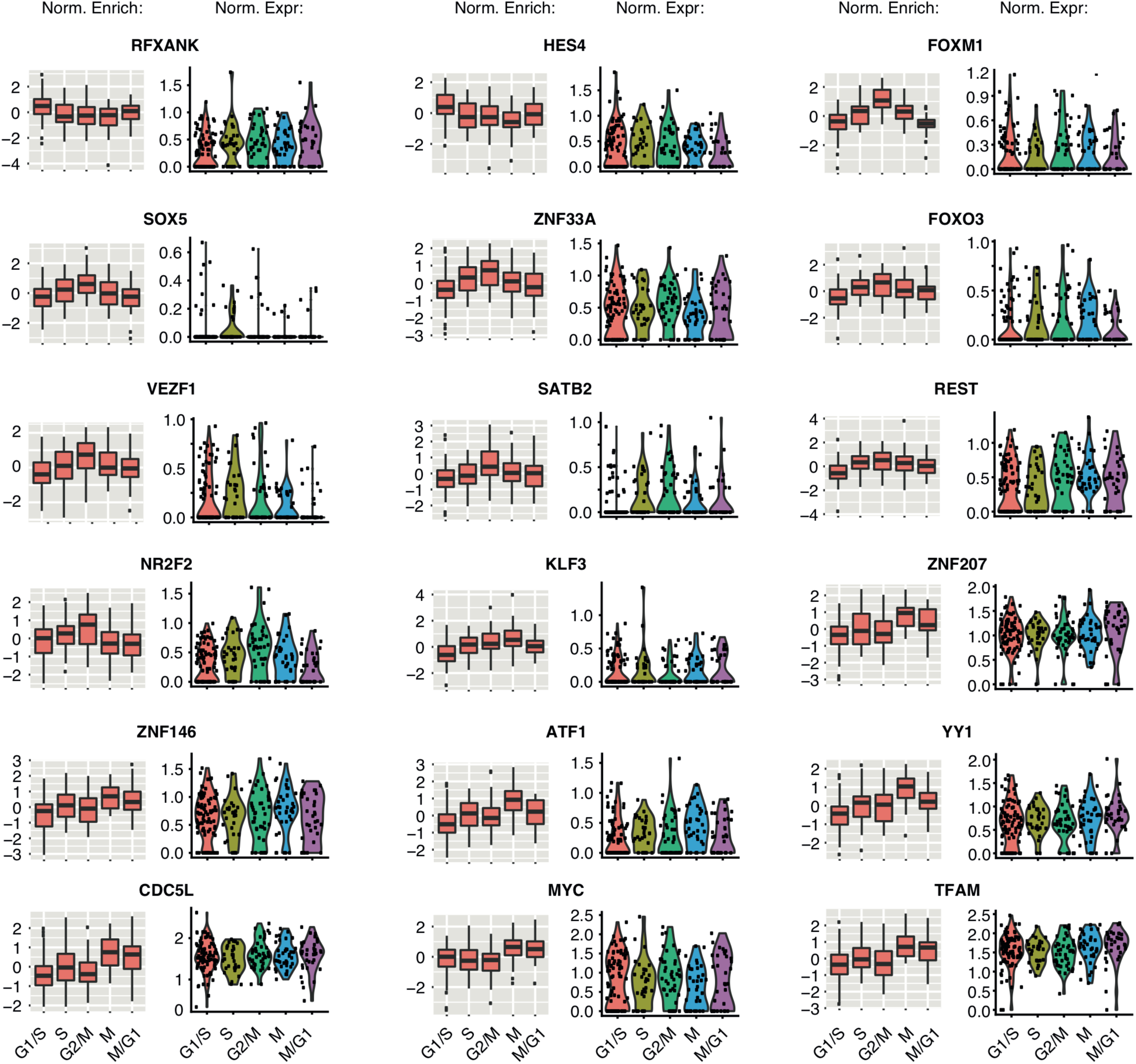
Boxplots showing normalised inferred activity and normalised gene expression across different phases for selective transcription factors shown in figure 3D. Normalised activity and expression scores for each transcription factor were mean centred across all cells before being summarised by assigned cell cycle phase.

